# Modeling the dynamics of progression of chromosomal alterations in cervical cancer: a computational model

**DOI:** 10.1101/121814

**Authors:** Augusto Cabrera-Becerril, Cruz Vargas-De-León, Sergio Hernández, Pedro Miramontes, Raúl Peralta

## Abstract

Computational modeling has been applied to simulate the heterogeneity of cancer behavior. The development of Cervical Cancer (CC) is a process in which the cell acquires dynamic behavior among non-deleterious and deleterious mutations, exhibiting chromosomal alterations as a manifestation of this dynamic. To further determine the progression of chromosomal alterations in precursor lesions and CC, we introduce a computational model to study the dynamic of deleterious and non-deleterious mutations as an outcome of tumor progression. Analysis of chromosomal alterations mediated by our model reveals that multiple deleterious mutations are more frequent in precursor lesions than in CC. Cells with lethal deleterious mutations would be eliminated, which would mitigate cancer progression; on the other hand, cells with non-deleterious mutations would become dominant, which could predispose to cancer progression. The study of somatic alterations by computer simulations during cancer progression provides a feasible pathway for insights into the transformation of cell mechanisms in humans. During cancer progression, tumors may acquire new phenotype traits, such as the ability to invade and metastasize or to become clinically important when they develop drug resistance. Chromosomal alterations non deleterious contributes to this progression.

## 1 Introduction

The genome and chromosomal unbalance of a transformed cell is highly heterogeneous, with a wide range of structural and copy number alterations. On the other hand, the behavior of the tumor as a whole, results from the accumulation of all of altered cells [1]. In particular, in Squamous Cell Carcinomas (SCC), there is strong experimental evidence for copy-number unbalance correlating with tumor aggressiveness [2]. In this regard, a large number of chromosomal alterations could characterize an aggressive tumor, while a smaller number of these alterations could be associated with a less aggressive tumor [3]. Therefore, finding the progression of these alterations would allow to address the cancer treatment based on the profile of chromosomal alterations.

On the other hand, although all number cells can undergo chromosomal alterations, only a small have the potential to be deleterious to the cell, while the majority of chromosomal alterations being not deleterious [4]. In this context, the development of cancer is result of the accumulation of several non deleterious mutations. In early stages of cancer, non-deleterious alterations are few, however, in advanced stages these alterations are considerable [5]. Several studies in molecular cytogenetics (e.g., comparative genomic hybridization) have investigated chromosomal alterations in cancer. Some of the authors of these studies correlate these chromosomal alterations with specific tumor behaviors[6]-[9].

Cervical Cancer (CC) is the second most common malignancy in women worldwide. The infection with high-risk Human PapillomaVirus (hr HPV) is the major etiological factor for this tumor [10]. Hr HPV is able to transform the infected cells by the direct action of the products of two of its early genes: the E6 and E7. E6 and E7 proteins of hr HPV are able to interact with molecules important for growth regulation and cell replication, as well as for repairing damage to the DNA of healthy cells. The E6 protein of hr HPV binds with high affinity to the molecule known as p53, inducing its degradation. The p53 protein is an important regulator of cell replication and is known as the main tumor repressor in humans; p53 is able to detect damage to the DNA and to arrest cell replication. A high proportion of human cancers have shown to have damage in the gene encoding the p53 protein; CC comprises an exception because, in this case, the gene is intact, but the protein is not present in cells infected by hr HPV, and E6 has been responsible for removing it. In this case, the cell cannot repair DNA errors and experience tumor development when the number of mutations increase. On the other hand, the E7 protein binds specifically to the tumor repressor gene product Rb. This gene was discovered and characterized in retinoblastoma [11]. This is a cell-cycle regulatory factor and is directly linked with the E2F transcription factor that, in turn, induces the transcription of elements involved in cell replication. E7 protein of hr HPV possesses high affinity for Rb-E2F. When E7 protein binds to Rb, E2F is released and induces cell proliferation. Thus, E6 and E7 cooperate efficiently in cell transformation, stimulating chromosomal alterations in the uterine cervix. The profile of chromosomal alterations is very heterogeneous in precursor lessions (cervical intraepithelial neoplasms 1, 2, and 3), but this heterogeneity decreases in CC. In advanced tumors, we observe a profile of chromosomal alterations induced by HPV oncoproteins [11, 12]. Using computational tools, it is possible to obtain a model of progression of chromosomal alterations since precursor lessions until CC[13]-[19]. Within the context of a cancer, these models help in determining global behavior of the tumor [20, 21]. The aim of this study is to determine the progression of chromosomal alterations such as deleterious or non-deleterious alterations, from precursor lessions to CC, using a computational model.

## 2 Computational model of cervical cancer

The molecular biology methods applied to study chromosomal alterations in CC indicate a great heterogeneity of such alterations. Thus, CC behaves as a complex system, rendering computational tools ideal for the study of the behavior of this tumor.

Agent-Based Modeling (ABM) is a computational modeling technique for the study of complex systems, i.e., systems that are composed of many interacting elements.The main idea of ABM is to replicate, with a stimulus, some of the interactions among individual components of the system.The ABM consists of the following three main components: agents, rules that govern interactions among agents, and the environment where agents interact. Currently, there are many options and platforms for developing ABM; however, its application in the study of cancer is recent [26, 24, 25]. For example, in a biological system such as tissue, each cell can be represented as an agent, and these agents may receive signals and input from environment and their neighboring agents, providing output to the environment and their neighbors and making decisions based on the input from around them and from their internal signals. Within this context, the dynamic of chromosomal alterations, from precursor lessions to CC can be modeled by an ABM. Under conditions in which a deleterious mutation is present, cells enter into an apoptotic state. If a non-deleterious mutation is active, the cell progresses to the development of advanced lessions and cancer. There are at least three advantages of utilizing ABM as a mechanism of modeling cellular cancer: multi-scale modeling; randomness, and emerging behavior.Also, we can employ random distribution to simulate external stimuli and to take into account the stochastic effects, alway present in biological phenomena.

On the other hand, with Cellular Automata (CA) is possible make idealizations of physical systems, they are discrete dynamical systems both in time and in space. For example, a single cellular automaton consists of a line of cells, each with a value of 0 or of 1 (true or false). These values are updated in a sequence of discrete time steps, according to a definite, fixed rule [14]. CA produce complex behavior even with the simplest defining rules. In general, cells have a finite number *k* of possible values and may be arranged on a regular lattice in any number of dimensions. Some defining characteristics of CA are: discrete in space and in time; discrete states; homogeneous; synchronous updating. The rule of each cell depends solely on the values of a local neighborhood of cells around it, and its state depends only on its values in preceding steps. Many biological systems have been modeled by CA [15, 16]. The development of structure and patterns in the growth of organisms often appears to be governed by very simple and local rules; therefore, they are well-described by CA. The advantages of using CA for modeling is may be considered as parallel processing computers as well; thus, complex behavior that involves many individual cells, may be modeled properly with CA. Cervical cancer has been modeled in different works using CA combined with another techniques [17]. In this work we model cellular behavior using CA and cellullar dynamics with an ABM, to determine the dynamics of deleterious and non-deleterious alteration in cervical tissue, therefore, our agents are autonomous, probabilistic-state cellular automata.

## 3 Overview of the model

The “ABM-Cervical-Cancer” model is a hybrid, two-Dimensional (2D) computational model implemented on the Netlogo framework [18]. It consists of a set of agents that simulate cell behavior. We decided to construct each cell with a minimal chromosome that is subjected to two possible alterations, deleterious and non-deleterious mutations, and each gene has three possible states: silenced; normal, or overexpressed. **Figure 1**.

**Figure 1:**
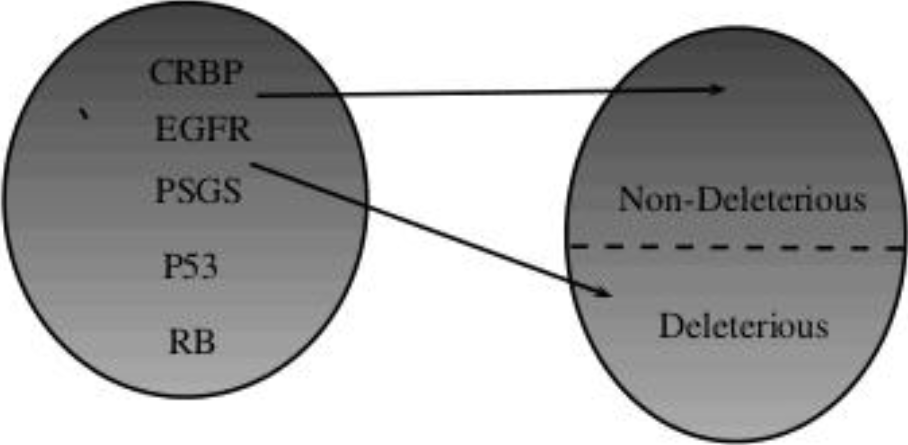
Minimal chromosome alterations in a cell.

We introduce a global variable “HPV” as a Boolean variable. When HPV is true, it means that there is infection with HPV, integration in genome, and the triggering of random alterations, these behaviors suggested by the literature [30].

The variable *P* simulates the clinical manifestation of HPV infection, *P* is the outcome of the roll of a pair of dice: if the result is greater or equal than 2 (*P* ≥ 2) then, the HPV-infected host will probably exhibit a Cervical Intraepithelial Lesion (CIN)1 lession; if the result of the roll of the dice is greater or equal than 5 (*P* ≥ 5) it is more probable that a CIN1 lession evolve into a CIN2 lession, and if *P* ≠ 7 it is highly probable for the host evolve to a CIN3 lession and and for it, once in this state, to develop Cervical Cancer (CC) with probability *η*.

The evolution from HPV infection to CC is simulated by counting the number of cells that present early-stage cervical intra-neoplasic lession, a second roll of the dice yields the probability for progressing from CIN1 to CIN2, although some of the cells with CIN1 lession will probably recover with probability *β* = 1%. If some deleterious alterations are present, death cell will be more probable and cancer will not develop. The “natural” cell-death rate is *μ*. Individual cells have two forms of interaction. Each cell breeds a new cell when it reaches some “mature” stage (when it passes a fixed time *τ*), the breed will inherit the chromosome of the “parent-cell”. Even more so, when a large number of cells neighbors are cancerous, the probability of transition from CIN2 to CIN3 or cancer is increased (**Figure 3**). In the **Appendix**, we have include a detailed description of the model using the ODD (Overview, Design concepts and Details) protocol.

**Figure 2:**
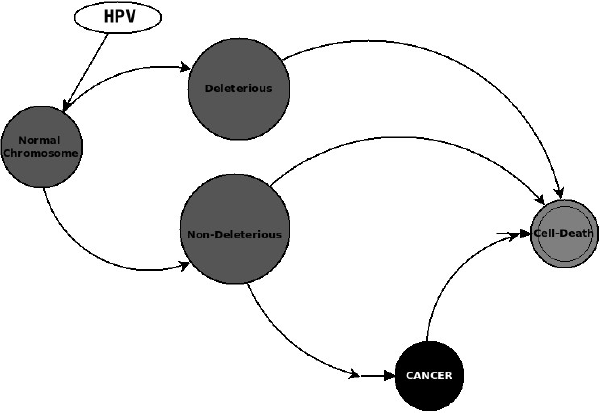
Chromosomal alteration dynamics.

**Figure 3:**
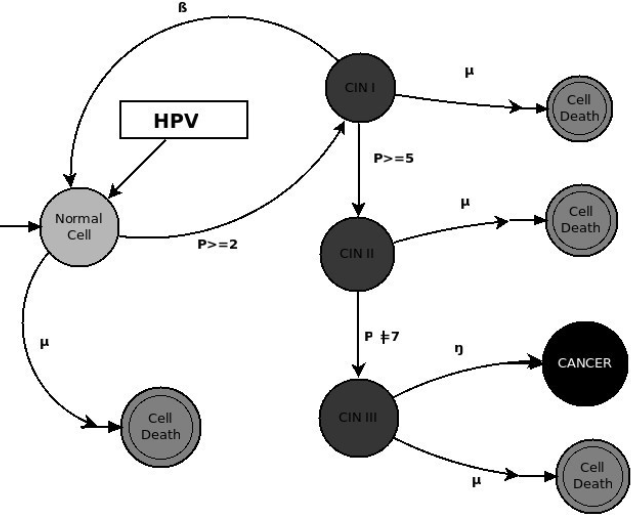
The ABM-Cervical-Cancer model.

In our model, the Clinical Intervention variable is an external stimulus that randomly selects a patch of cancer cells and deletes it in one step of the simulation; this allows us to simulate real clinical intervention. In the next step, the dynamics repeat; thus, cancer cells remain present after Clinical Intervention; their growth is being bound. The mechanism produces loops in the simulation, as depicted in **(Figure 2 and 4).** Additionally, the randomness produces oscillation in the dynamics of the model.

**Figure 4:**
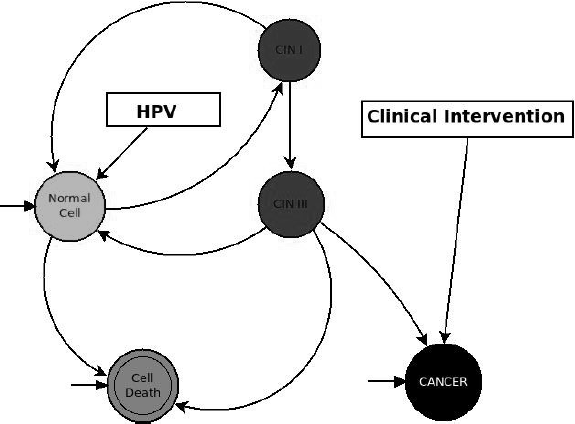
Dynamic of ABM-Cervical-Cancer model with Clinical Intervention variable.

The cells possess two ways of interaction: “horizontal” interaction and “vertical” interaction. Vertical Interaction refers to the inheritance of characteristics or mitotic transmission. Horizontal interaction occurs with the nearest cells in a De Moore neighborhood. Horizontal Interaction is governed by a random process; thus, the probability for a single cell to develop cancer increases when cancer cells are nearby in its neighborhood **(Figures 5, 6 and 7).**

**Figure 5:**
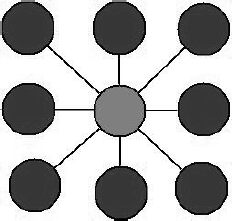
Neighborhood for ABM-Cervical-Cancer model.

**Figure 6:**
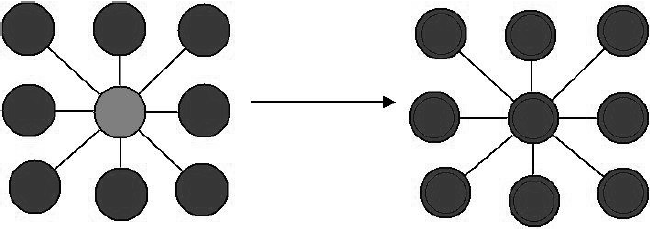
“Horizontal” Interaction.

**Figure 7:**
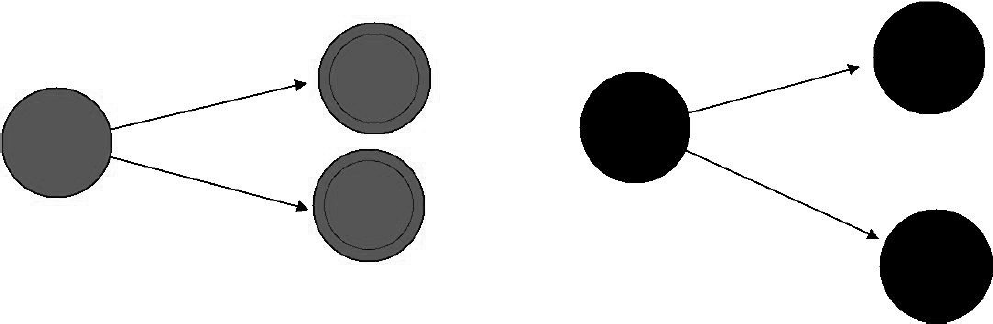
“Vertical” Interaction or mitotic transmission.

The condition for halting the program consists either of having a 75% cancer cells spread on the tissue, or else reaching 20,000 steps of the simulation. It is noteworthy that in this model, we do not pretend to simulate the growth of cancer mass, but instead, to simulate chromosomal variation as the cancer cells growth. Therefore, the movement of the agents does not correspond to the natural dynamics of tissue growth, but chromosomal alterations are rather simulated carefully.

## 4 Simulation and Results

We define two different experiments, each consists on 100 runs of 20,000 iteration of ABM-Cancer on the NetLogo framework, fixing some initial conditions for the density of cell-growth and cell death-rate. For the initial experiments *E*_1_ and *E*_2_, Clinical Intervention has been set as false. For experiments *E*_1_1__ and *E*_2_1__ has been set as true. The main difference between the *E*_1_ and *E*_2_ is the election for cell density and death rate, as shown in the Table 1.

**Table 1:** ABM-Cervical-Cancer initial setup for numerical experiments

| Experiment | HPV | Cell count | Cell death-rate | Clinical Intervention |
| --- | --- | --- | --- | --- |
| $E_1$ | true | 1740 | 0.01 | false |
| $E_2$ | true | 6000 | 0.02 | false |
| $E_{1_1}$ | true | 1740 | 0.01 | true |
| $E_{2_1}$ | true | 6000 | 0.02 | true |

Experiments *E*_1_ and *E*_2_ have shown similar statistical behavior. In 18% of runnings of experiment *E*_1_ CIN1 lessions appear, which is consistent with 16% prevalence of cervical cancer due to HPV infection, as reported in the literature [23, 22]. Experiment *E*_2_ reveals a 12% prevalence of CIN1 lessions. Both experiments showed a different mean duration, while nearly all of the CIN1 positive cases of *E*_1_ had invasive cancer, in experiment *E*_2_ runs almost none of them presented invasive cancer (Table 1).

On the other hand, runnings in which Clinical Intervention was set as true present a minor prevalence of CIN1 lessions as expected. In experiment *E*_1_1__ there was 11% prevalence, but in experiment *E*_2_1__ we obtain a 12% prevalence, the same as in *E*_2_ with the absence of Clinical Intervention. Experiment *E*_2_1__ present interesting behavior: although Clinical Intervention is present, in cases when CIN1 lessions appear, invasive cervical cancer is more likely to develop; therefore simulation does not reach the 20,000 steps, which may be a pattern caused by reaching the charge capacity of the system.

In Table 2 we show the arithmetical means of results on numerical experiments of ABM-Cervical-Cancer, which summarizes the statistical behavior of the model.

**Table 2:** ABM-Cervical-Cancer statistical results from numerical experiments

| Experiment | Max Steps | CIN1 prevalence | deleterious | non deleterious | Ratios |
| --- | --- | --- | --- | --- | --- |
| $E_1$ | 6400 | 18% | 297.18 | 317.12 | 1.03 |
| $E_2$ | 20000 | 12% | 250.23 | 264.35 | 1.05 |
| $E_{1_1}$ | 20000 | 11% | 123.83 | 148.34 | 1.2 |
| $E_{2_1}$ | 18 | 12% | 86.5 | 102.5 | 1.1 |

Dispersion analysis shows that there is a correlation between heterogeneity of chromosomal alterations and cancer progression. Analysis of time series show that in cases where CC has developed non-deleterious transformed cells are more prevalent that deleterious transformed cells. In subsequent figures we depict some runs of experiment *E*_1_. **(Figure 8.)**

**Figure 8:**
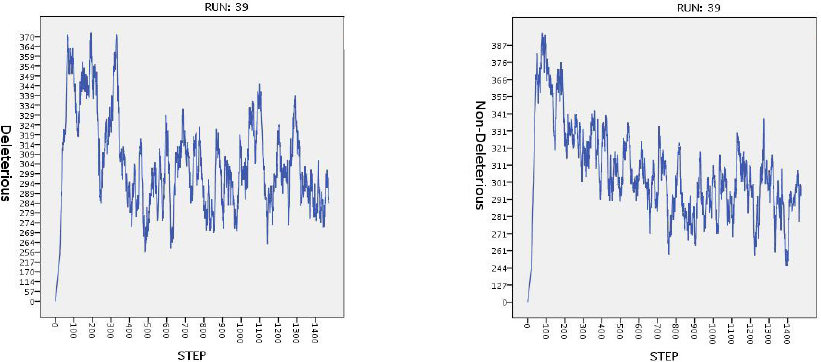
Time series for run 39 of experiment *E*_1_

In experiments *E*_2_ and *E*_1_1__, simulations reach the 20,000 steps and in both, we can observe that oscillation occurs in both deleterious and non-deleterious alterations. In experiment *E*_1_ we have a more interesting behavior: we have noisy oscillation in both alterations, although in the case of *E*_1_1__, oscillations are bound due to Clinical Intervention. On the other hand, *E_2_* exhibits an initial spike in both alterations, produced by the fast spread of the cancer cells, but eventually, the number of transformed cells reaches a stationary state. The main difference among these three is that only in experiment *E_1_* we were able to reproduce invasive cancer. Therefore, it is in this experiment that we expected to find the more interesting behavior.

In **(Figure 9)** we present the histograms of frequencies and recounts of deleterious an non deleterious alterations for run 8 of *E*_1_1__.

**Figure 9:**
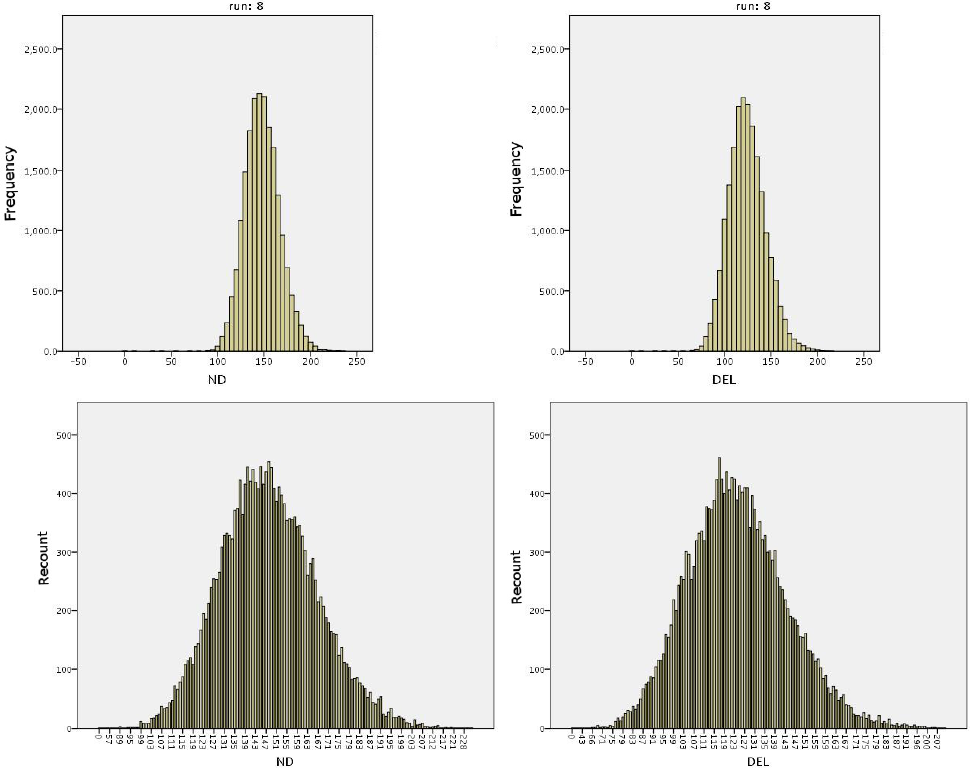
Histograms of frequencies and recounts in run 8 of experiment *E*_1_1__. ND stands for non deleterious and DEL stands for deleterious.

Progression of chromosomal alteration when non-deleterious mutation is present, will probably drive to CC, acting as an selective mechanism; but when deleterious mutations are present, cell death will probably arise, this can be observed in the results of our simulation. In the figure below we illustrate the progression of CIN1 and CIN3 lessions and the progression of cervical cancer in run 39 of experiment *E*_1_ (we choose arbitrarily run 39).**(Figure 10)**

**Figure 10:**
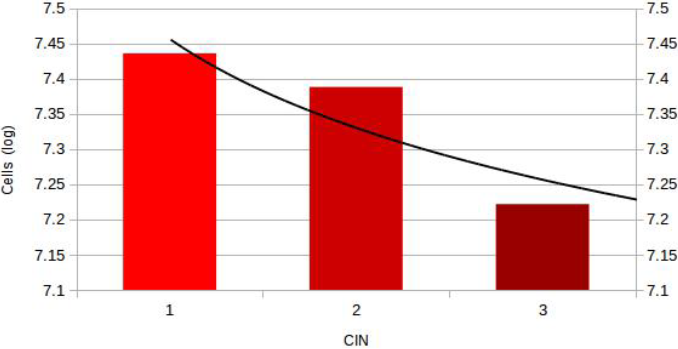
CIN lession progression with deleterious alteration.

## 5 Discussion

The development of cancer can be seen as a process of accumulation of mutations and chromosomal alterations, with the selection of cells that, to the best of their ability, adapt to their perturbed environment. Survival of the transformed cells is driven by the sequential accumulation of genetic alterations in sets of genes that control cell proliferation and the diferentiation of signal transduction pathways. The number and types of somatic mutations accumulated during cancer progression are variable among different cancer types. Alterations typically include allele loss, point mutations, amplifications, and others, but these alterations are essentially divided between deleterious and non-deleterious to the cell [27].

In the human genome, there are specific regions that can potentially regulate cellular behavior through their control of numerous intra- and intercellular signaling networks. Some genome products (non-coding RNA, oncogenes, tumor suppressor) already comply with the criteria of being important nodes; however, understanding how they influence different pathways is limited by the lack of knowing how they participate in different complexes and networks and how their mutation-level status affects complex composition and functions within cell types in tumor tissues. Genetic and epigenetic networks are altered in cancer cells, and they reflect permanent changes in the types and interactions of network complexes at several levels, including the cell cycle and differentiation of signal transduction pathways, in which their alterations could have deleterious effects to the cell. Deleterious regions may be important hubs that make multiple (functional or physical) connections with nodes that control tumor-cell behavior. It is currently not clear how the products of deleterious regions are organized into networks. However, recent evidence suggests that they may be organized as critical hubs in clusters among networks, with some of these mutated in the majority of cancers [28, 30].

Some mutations are potentially deleterious to the cell, and in cancer progression, these do not accumulate. A cell with mutations, could have both desirable (non-deleterious) and undesirable (deleterious) consequences. Cells with lethal deleterious mutations would be eliminated, which would mitigate cancer progression. However, cells with non-deleterious mutations would become dominant, which could predispose to cancer progression. During cancer progression, it becomes increasingly harder to eliminate deleterious mutations or to fix beneficial mutations [31]. The number of documented somatic copy number alterations are variable, but random mutations may help explain why neutral passenger mutations (non-deleterious) are common in carcinomas. Interestingly, cancer genome mutation frequencies are related with cancer progression, suggesting that many non-deleterious mutations first accumulate in transformed cells. The ratio of non-deleterious to deleterious mutations (ndm/dm) is essentially the expected value of random mutation, suggesting that the majority of non-deleterious mutations in cancer genomes were accumulated. Interestingly, non-intuitive phenomena frequently emerge during cancer progression. Multiple deleterious mutations are most frequently found in the precursor lession, which is invasive cancer [32]. Given the unfeasibility of human experimental manipulations, the analysis of somatic alterations by computer simulations during cancer progression provides a feasible pathway for insights into the human transformation of cell mechanisms. Somatic alterations can reveal much about the cancer cell during progression. During cancer progression, tumors may acquire new phenotype traits, such as the ability to invade and metastasize, or may become clinically important when they develop drug resistance. Acquired chromosomal alterations contribute to this progression [33].

In the cervical epithelium, virtually all premalignant lessions and cancers are caused by HPV, and chromosomal instabilities could be linked with the effect of deregulated E6 and E7 expression. Clinically, epithelial lessions associated with transforming HPV infections demonstrate a heterogeneous course, with only a proportion of high-grade lessions progressing to invasive cancer; thus, biologically, progression toward cancer is likely the consequence of the accumulation of specific and multiple cellular changes that promote the outgrowth of distinct cell clones. The risk for the progression of a premalignant lession to cervical cancer is further linked with the HPV genotype: infection with HPV 16 and 18 are substantially more likely to progress in less time in comparison with lessions induced by other high-risk HPV types [17]. The introduction in our model of variable specific HPV could potentially modulate the progression of precursor lessions to CC.

Computational models such as Cellular Automata (CA), Agent Based Models (ABM), and their hybrids are techniques in silico for studying a variety of cancer behavior. Using hybrid computational techniques, it is possible to simulate the role of diversity in cell populations as well as within each individual cell. Therefore, it has become a powerful modeling method that is widely used by computational cancer researchers. Many aspects of cancer progression, including phenotype-changing mutations, adaptation to the microenvironment, the process of angiogenesis, the influence of extracellular matrix, reactions to chemotherapy, the effects of oxygen and nutrient availability, and metastasis and the invasion of healthy tissues have been incorporated into and investigated in computational models [34, 36].

In a hybrid model, each cell is often represented as an agent that behaves locally as a CA. Agents may receive signals and input from the environment and from their neighboring agents and make decisions based on the input surrounding them and their internal environment. Within this context, an agent may grow, proliferate, or undergo apoptosis in response to the surrounding environmental conditions [37]. During cancer progression, cellular proliferation requires genomic stability; if not, it will undergo apoptosis. In this regard, the dynamics of deleterious and non- deleterious mutations can be simulated as a manifestation of cell death mediated by apoptosis; i.e., deleterious mutations in precursor lessions are lethal to the cells. Thus, the latter do not progress to advanced lessions and cancer, while non-deleterious mutations in precursor lessions, induce cellular transformations and cancer, rendering hybrid modeling an ideal tool to model this process.

## 6 Conclusions

In this paper, we focused our eforts on the progression of chromosomal alterations in cervical cancer employing a hybrid computational model, especially on the manner in which deleterious and non-deleterious alterations affect cancer behavior across different precursor lessions in CC.

Interestingly, this study revealed significant high frequency in deleterious mutations in precursor lessions and low frequency in these mutations in CC. Precursor lessions of the cervix are well defined and its progression to CC shows that can be seen as an accumulation of deleterious mutations. Deleterious mutations are more common in precursor lessions, but these do not occur in later or advanced stages of cancer.

Some extensions for the model may include new variables, for instance, a vaccination variable. Vaccination could be simulated as another external stimulus that modulates the clinical manifestation of HPV infection; therefore, it would be a weight function of the P variable. We can also extend the model to simulate interactions among scales; thus, we can begin to elucidate how chromosomal alteration links up with other phenomenological manifestations of cervical cancer, fabricating a multi-scale version of the ABM-Cervical-Cancer model.

## Appendix: ODD for ABM-Cervical-Cancer

- Overview In some cases, the apparition of cervical cancer (CC) is preceded by three types of sequenced lessions (CIN1, CIN2 and CIN3) after the infection of HPV virus. The infection of HPV can produce chromosomal alterations. Progression of chromosomal alterations in cervical cancer has the main purpose of showing how different alterations, simulated by different values of random variables as external stimuli over the selected chromosome and the action of medical stimuli as well, can induce cervical cancer
- State Variables and Scales

– Chromosome: Is an ordered pair (x,y) that will simulate the affected chromosome, it that reflects each of the stages in the model, x will reflect the appearance of cervical cancer (CC) and y will reflect the appearance of cervical cancer. The values that x can take models the different lessions that are associated previous to the appearance of CC and determined by a random variable, and another random variable will govern the state of y, that is either cancer is present or not. HPV is a Boolean variable that when set to True means that an infection of HPV on the tissue that can trigger random alterations, False otherwise
– Clinical Intervention is a Boolean variable that when set to True, altered tissue cells are removed physically
– *P* is a variable that simulates the strength of the host immune system. It is a result of a double dice rolling where the random variable (*X*) is the sum of the “visible” faces of the dice. If *X* ≥ 2 then the host is infected with HPV and and then in a Bernoulli process we will determine if it does gets infected. CIN1 lessions will appear if *X* ≥ 5 then it is more probable that CIN1 will become a CIN2 and also if *X* ≠ 7 then it is highly probable to evolve to a CIN3 lession and from this state develop Cervical Cancer.
- Process Overview and Scheduling The evolution of HPV to CC is simulated counting the number of cells that present early stage and then another dice rolling will determine the probability of transition from CIN1 to CIN2. Figure 2 shows the schedule and transition of the model. In figure 3 it is shown the gene alteration dynamics that occurs when the cell is infected with HPV. The figure shows the transit to apoptosis when deleterious mutations occurs, but when non-deleterious mutations occurs we can have the proliferation of cancerous cells. In figure 4 it is shown the model when the Clinical Intervention variable is set to true and how the removal of cancerous cells will occur at different stages of the model.
- Design Concepts Since this model is thought to examine the relationship between the presence of CC given a different set of lessions and how this CC evolves in the presence of Clinical Intervention, the design concepts required to be mentioned to this regard are as follows. A double random process is required in order to pass from a specific type of lession to another, when the chromosome can present some type of lession then another random variable will determine if this lession (or CC) will be presented or not. Some of the cells with CIN1 lession will probably recover with probability of 1%. If deleterious alteration are overexpressed, death cell will be more probable and cancer will not develop. Clinical Intervention is an external stimuli that randomly chooses a number of cancer cells an deletes them, and the process is repeated in a next step
- Details
  – Initialization The initialization is set with HPV false for all chromosomes and it will run over 100 runs over 20,000 iterations with the Clinical Intervention variable set to false, another experiment is made with Clinical Intervention variable set to True.
  – Input The model does not require specific input other than initialization values
  – Submodels There are no particular submodels used in ABM-CC

## Conflict of interests

The authors declare that they have no conflict of interest.

## Acknowledgments

RP wishes to thank SEP-PRODEP 6986 for the support. The authors also wish to thank to Dr. Mauricio Salcedo for their constructive comments and suggestions.

